# A fast, robust method for quantitative assessment of collagen fibril architecture from transmission electron micrographs

**DOI:** 10.1101/2023.02.06.527383

**Authors:** Bruno V. Rego, Dar Weiss, Jay D. Humphrey

**Affiliations:** Department of Biomedical Engineering School of Engineering & Applied Science Yale University, New Haven, CT, USA; Vascular Biology and Therapeutics Program Yale School of Medicine, New Haven, CT, USA

## Abstract

Collagen is the most abundant protein in mammals; it exhibits a hierarchical organization and provides structural support to a wide range of soft tissues, including blood vessels. The architecture of collagen fibrils dictates vascular stiffness and strength, and changes therein can contribute to disease progression. While transmission electron microscopy (TEM) is routinely used to examine collagen fibrils under normal and pathological conditions, computational tools that enable fast and minimally subjective quantitative assessment remain lacking. In the present study, we describe a novel semi-automated image processing and statistical modeling pipeline for segmenting individual collagen fibrils from TEM images and quantifying key metrics of interest, including fibril crosssectional area and aspect ratio. For validation, we show illustrative results for adventitial collagen in the thoracic aorta from three different mouse models.

## 1. INTRODUCTION

Collagen, the most abundant protein in mammals, provides critical mechanical support and strength to a wide range of soft tissues, including cartilage, ligaments, skin, and vascular tissues [1]. The multiscale structure of collagen is complex, consisting of a hierarchical assembly of molecules, fibrils, fibers, and ultimately fiber bundles [2]. From a biomechanical perspective, the architectural characteristics are important at nano-, micro-, meso-, and macro-scales, giving rise to the all-important properties of the collagen network (e.g., fiber orientation and undulation distributions) that are most frequently considered in both empirical and modeling studies [3–9]. For instance, the diameter of collagen fibrils correlates directly with tensile stiffness and strength in connective tissues [10]. Within blood vessel walls specifically, the architecture of collagen fibrils, including their size, organization, alignment, and spatial density, has been shown to be tightly coupled to the tissue-level mechanical behavior and thus function [11–13]. Collagen fibril density, diameter, and cross-sectional shape, as well as cross-linking, also influence the development and progression of various vascular pathologies, such as aneurysms, dissections, and ruptures [14–19]. In addition to affecting mechanical integrity and function, fibril architecture plays a critical mechanobiological role in the vasculature. The mechanical loads acting on blood vessels, namely blood pressure-induced intramural stresses and flow-induced shear stresses, have been shown to regulate various cellular processes, including proliferation, differentiation, and apoptosis [20], and the collagen fibril architecture within the vessel wall modulates these mechanobiological responses through its impact on cell–matrix interactions [18, 21].

Experimentally, transmission electron microscopy (TEM) has proven to be a powerful technique to study the architecture of collagen fibrils in various tissues [12, 13, 18, 22]. The high resolution and contrast of TEM images enables detailed analysis of the cross-sectional geometry, spatial density, and organization of collagen fibrils, all of which relate directly to mechanical properties and mechanobiological metrics of interest. Despite the value of TEM as an imaging modality, however, there is a lack of computational tools for processing TEM images—each of which typically contains hundreds of collagen fibrils—to quickly, objectively, and quantitatively assess fibril architecture across experimental groups with moderate-to-high sample sizes (e.g., dozens). Existing approaches for TEM-based collagen fibril segmentation remain impractical for high-throughput applications since they rely heavily on manual tuning of image processing parameters and/or interactive tracing of the individual fibril boundaries [13]. While computational image analysis protocols have the potential to reduce processing times significantly, no fully automated algorithms have yet proven to be robust for architectures that involve substantially variable fibril cross-sectional geometries [22]. Since pathological tissues, in particular, often contain collagen fibrils that vary widely in terms of size, boundary shape, and distribution as well as their overall visual appearance in TEM images (e.g., non-uniform intensity) [11, 13], there is a pressing need to leverage and extend automatic image segmentation techniques (which enable high throughput) within an interactive software package that allows for user guidance and verification (to ensure accurate segmentation and analysis results).

In the present study, we developed a novel semi-automated pipeline to segment individual fibrils from TEM images. While based primarily on advanced image processing and statistical modeling algorithms that execute automatically, our approach is implemented within a graphical interface, thus allowing user supervision and interactive correction when necessary. Focusing on vascular applications, we validated our pipeline using TEM images of the thoracic aorta from both wild-type and diseased mice, whose collagen fibril characteristics vary considerably in terms of size, shape, spatial arrangement, and visual appearance.

## 2. METHODS

### 2.1. Illustrative data set

We applied our approach to three thoracic aorta specimens excised from adult (8–14 weeks of age) male mice having a C57BL/6 genetic background to demonstrate and validate our TEM image segmentation and analysis pipeline in the specific context of vascular applications. To test our methodology against substantially different collagen fibril architectures, we show results for one wild-type control ascending thoracic aorta (ATA), one descending thoracic aorta (DTA) with disrupted TGF-β signaling (denoted as *Tgfbr1r2*; see Kawamura *et al*. [13] for details), and one *Fbn1^C1041G/+^* ATA representative of Marfan syndrome (denoted as MFS; see Weiss *et al*. [23] for details). Specimen excision and preparation as well as image acquisition methods have been described previously [13, 23]. Briefly, segments were fixed in 2.5% glutaraldehyde/2% paraformaldehyde in sodium cacodylate buffer at room temperature for 2 h at 4°C. Samples were rinsed in sodium cadodylate buffer before post-fixation in 1% osmium tetraoxide for 1 h and subsequent staining using 2% uranyl acetate for 1 h. Afterward, samples were washed and embedded in resin and sections of the adventitial layer were imaged using a FEI Tecnai BioTwin transmission electron microscope.

### 2.2. Image processing and analysis

The fibril segmentation pipeline presented herein was implemented as a graphical user interface that combines automated image processing algorithms with periodic queries that allow the user to supervise and interactively correct intermediate outputs. This approach was chosen with the simultaneous objectives of maximizing efficiency while also minimizing errors in the final results. Below, we summarize step by step both the user involvement and the processing methodologies employed to analyze an individual image.

As an initial preprocessing step, the user interactively selects regions of the image to exclude from the analysis—for example, perivascular areas containing adipose tissue rather than collagen fibrils (Figure 1). The image is then smoothed using a Gaussian filter with bandwidth equal to 0.3% of the smallest dimension (height or width) of the image (Figure 2a,b). To regularize the intensity field of the image as a whole, a bivariate quadratic function

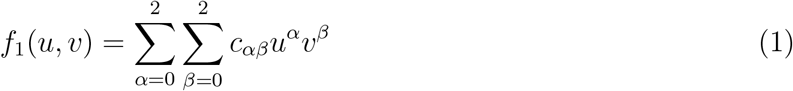

is first fit to the intensity values *I*_filt_(*u, v*), where (*u, v*) are pixel coordinates. To minimize the influence of spurious bright/dark pixels that deviate substantially from their average regional intensity, the coefficient values *c_αβ_* are optimized using robust iteratively reweighted least-squares regression with a Cauchy weight function,

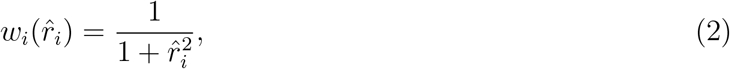

for

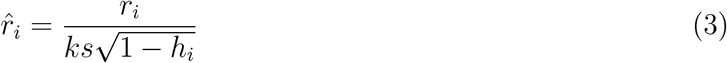

and

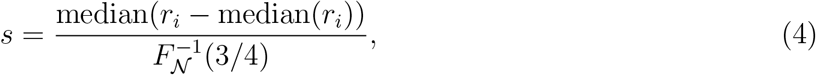

where *r_i_* are the ordinary least-squares residuals, *h_i_* are the least-squares fit leverage values, *s* is a robust estimator of the standard deviation of the residuals based on their median absolute deviation, 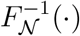 is the inverse of the standard normal cumulative distribution function, and *k* ≈ 2.3849 is a tuning constant whose value yields 95% asymptotic efficiency for normally distributed data [24].

**Figure 1:**
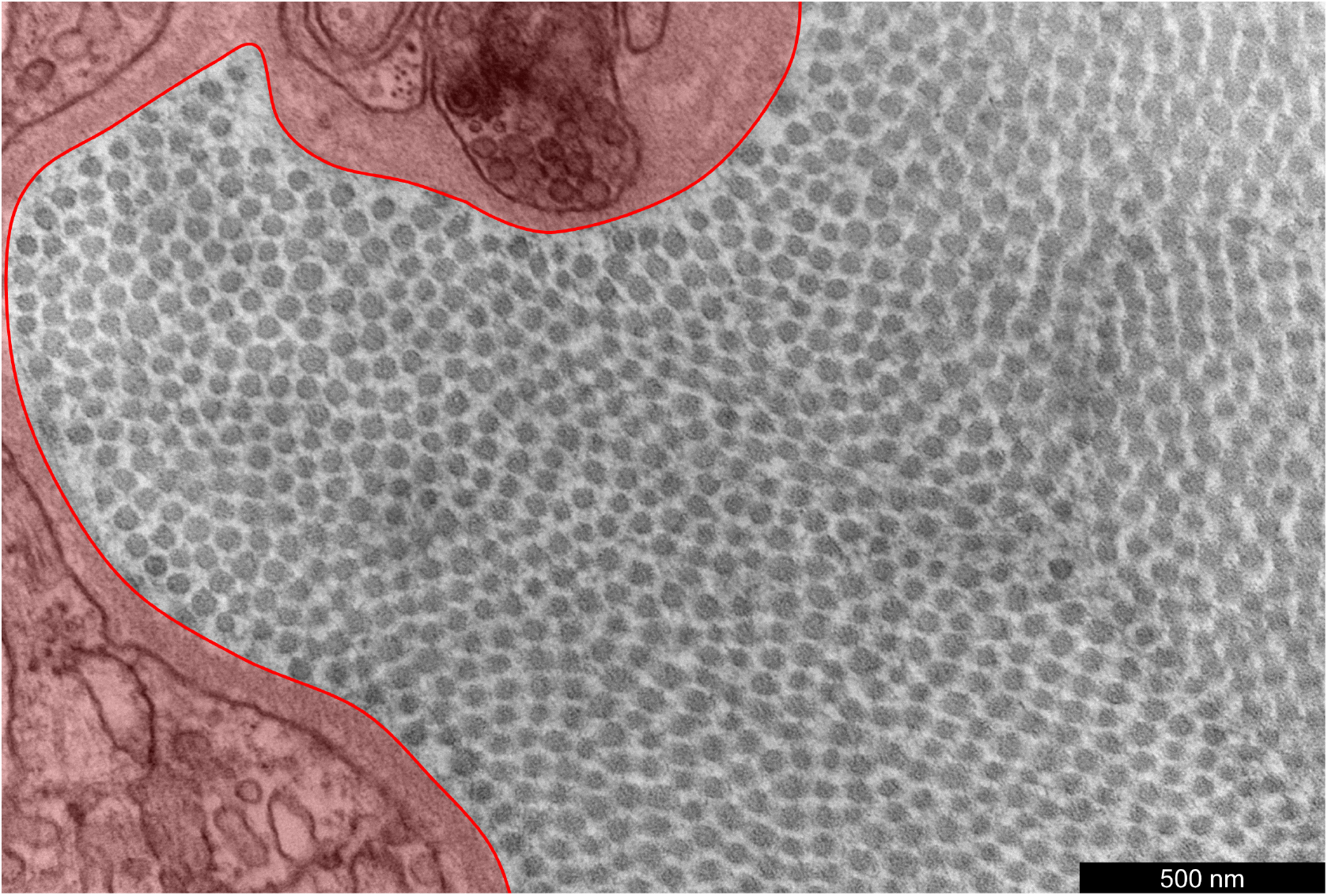
An illustrative TEM image of the adventitial portion of the ATA from a wild-type control mouse. The perivascular region highlighted in red has been interactively outlined to be excluded from analysis, since it does not contain collagen fibrils.

The modeled intensity values *f*_1_ (*u, v*) are adopted as a local adjustment factor, and the intensity values of the filtered image are adjusted using

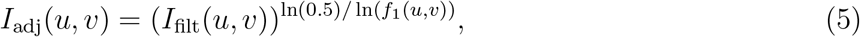

such that pixels with intensity *I*_filt_(*u, v*) = *f*_1_ (*u, v*) are mapped to 0.5. This spatially heterogeneous (yet smooth) adjustment attenuates the influence of regional variations in image intensity (e.g., due to non-uniform lighting) during the binarization step (Figure 2c). Using Otsu’s method [25], the transformed image is then binarized (*I*_bin_) according to an optimal threshold, with dark pixels categorized as fibrils (Figure 2d).

**Figure 2:**
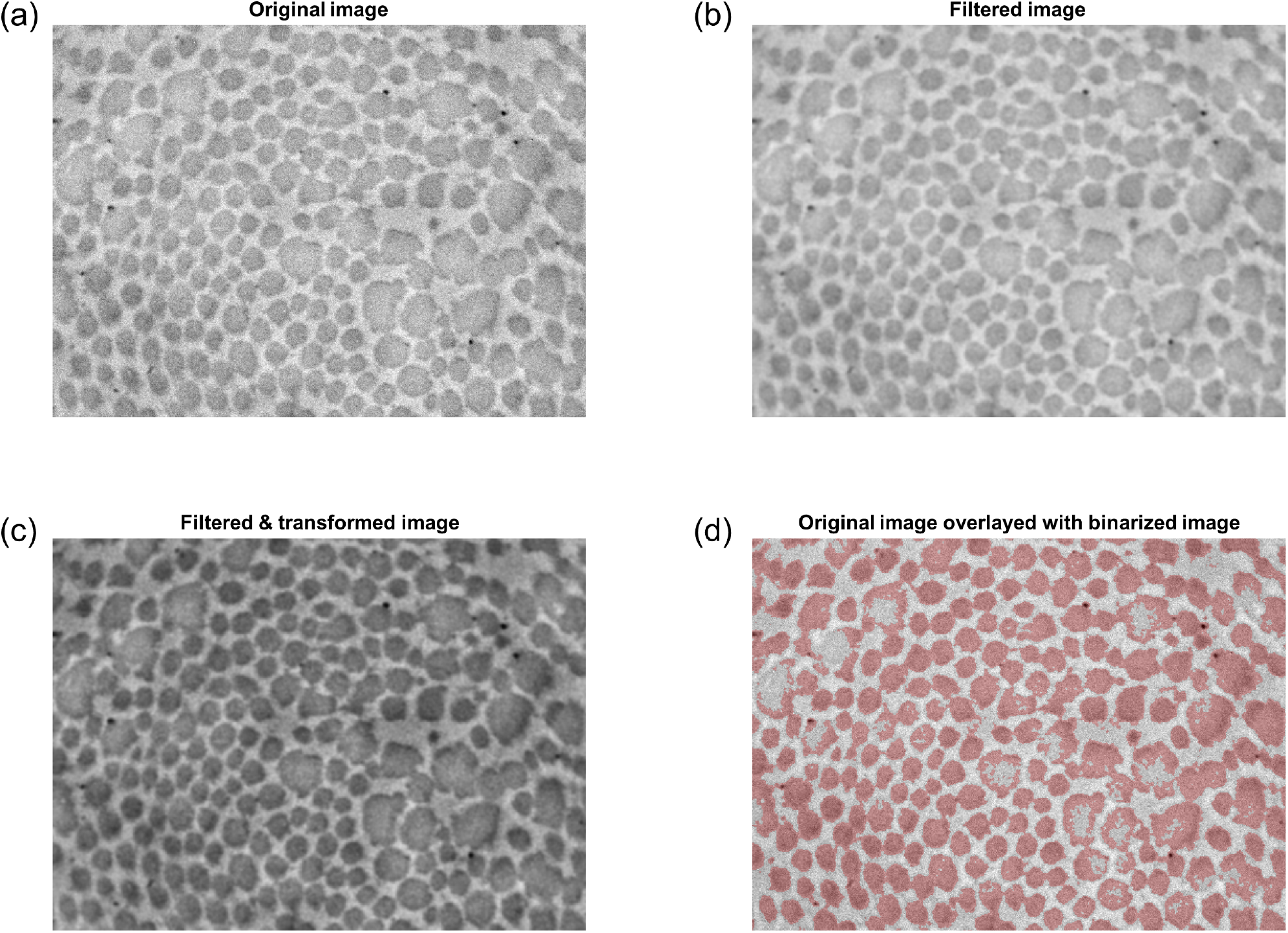
Filtering and binarization pipeline. (a) Original TEM image (*Tgfbr1r2* DTA). Note the moderate but perceptible variations in intensity across the image, with regions near the corners generally appearing darker than the middle. (b) Image after Gaussian filtering. (c) Image after subsequent power transformation via Equation 5, which attenuates regional variations in intensity. (d) Overlay of the original image (grayscale) with the binarized image (red). Note that in images of pathological vessels (such as the one shown), this first binarization attempt performs poorly in larger fibrils that have uneven intensity.

For each pixel in *I*_bin_, the Euclidean distance to the nearest non-fibrillar pixel is computed, producing a distance field *D* with the same dimensions as *I*_bin_. Then, for different values of the Gaussian filter bandwidth *σ_i_* (ranging from 0.5 to 10 pixels, in steps of 0.5 pixels), the number of local peaks *N*(*σ_i_*) in *D*_filt_ (*σ_i_*)—taken to be the number of “detected” fibrils—is computed (Figure 3a). A 5th-degree polynomial *p*(*σ*) is fit to the resulting (*σ_i_,N*(*σ_i_*)) data (Figure 3b), from which the optimal bandwidth *σ*\* is considered to be the lowest positive *σ* where (*dp/dσ*)^2^ reaches a local minimum (Figure 3c). This filter bandwidth corresponds to the lowest positive *σ* where the slope of *p*(*σ*) is locally the nearest to 0, meaning that the number of detected fibrils is minimally sensitive to the filter bandwidth at this point. Because (*dp/dσ*)^2^ is also a polynomial, it is straightforward to compute the roots of its derivative (one of which is *σ*\*), thus rendering iterative minimization of (*dp/dσ*)^2^ unnecessary.

**Figure 3:**
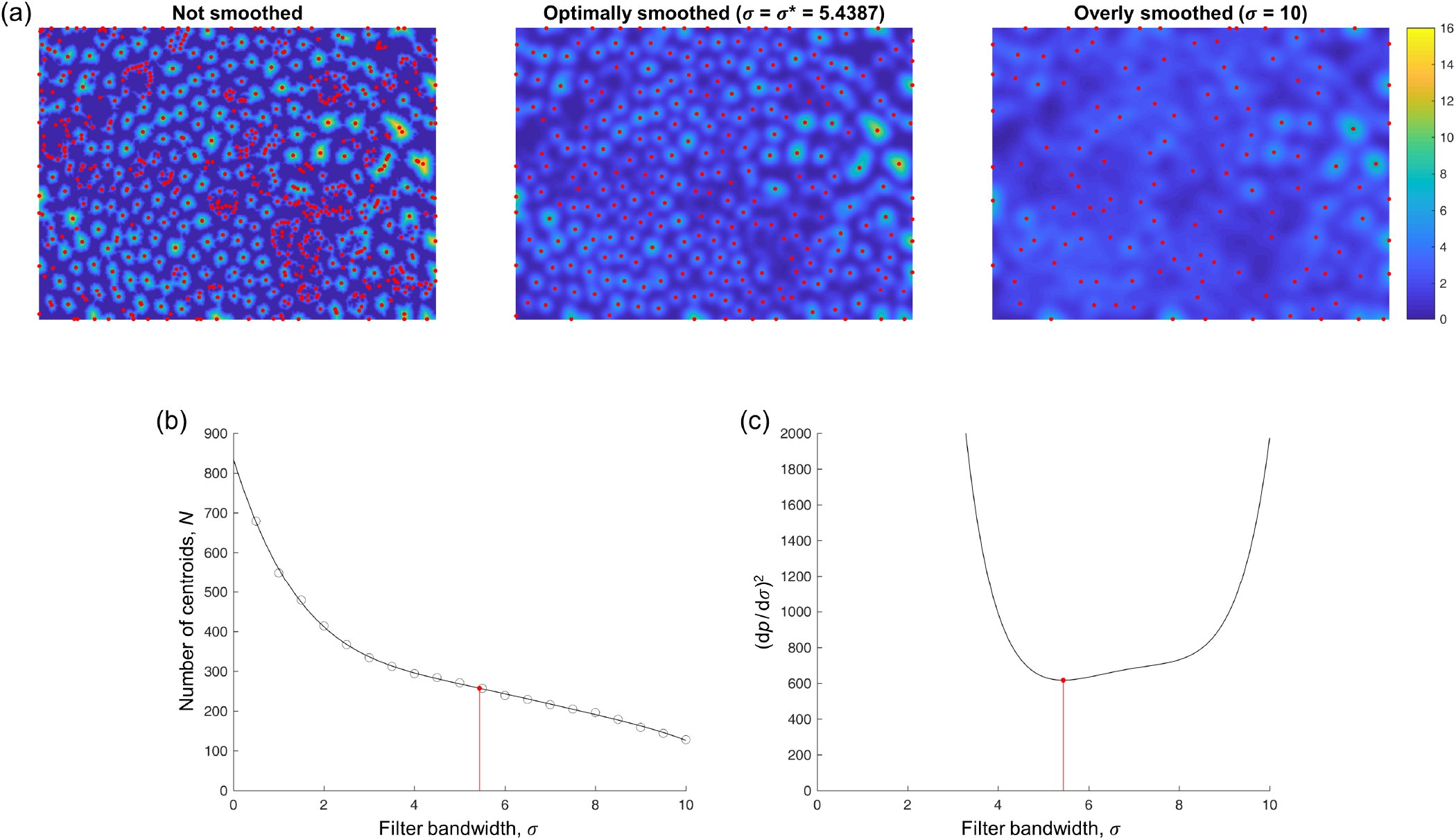
Fibril centroid detection. In (a), the color scale displays the distance field *D* (unsmoothed) or *D*_filt_ (smoothed with the indicated bandwidths *σ*). Local peaks in the field, plotted in red, are treated as candidate locations for the centroids of all the collagen fibrils in the image. (b) The number of centroids *N* detected with different filter bandwidths *σ*. Circles denote actual results, while the plotted curve corresponds to the 5th-degree polynomial fit *p*(*σ*). The optimal filter bandwidth (red) corresponds to the point at which N is least sensitive (locally) to *σ*. (c) Equivalently, the optimal filter bandwidth corresponds to the lowest positive *σ* where (*dp/dσ*)^2^ reaches a local minimum, which is straightforward to compute.

The first estimates of the centroid coordinates are thus the locations of the local peaks in Dfiit (*σ*\*), with the corresponding Voronoi tessellation defining a first estimate of the fibril “neigh-borhoods” (Figure 4a). At this stage, the user may interactively add, remove, or modify fibril centroids that were undetected, erroneously detected, or poorly estimated, respectively, if needed (Figure 4b,c). For each Voronoi cell, if any of the cell’s vertices are either outside of the image area or within a previously excluded region, the corresponding fibril is labeled as a “boundary fibril,” to be excluded from future analyses (e.g., cross-sectional area computations) since part of the fibril is mostly likely not visible. While small fibrils tend to have a uniformly dark intensity, the central region of relatively large fibrils (e.g., Figure 2a) is often lighter than the outer regions (and sometimes as light as the background). In these cases, the first binarization attempt tends to incorrectly classify the light central regions as non-fibrillar, thus necessitating a refinement to the binarization. To improve upon the initial image binarization, a “characteristic intensity” is defined for each Voronoi cell, equal to the mean intensity of pixels in that cell whose pixel-to-centroid distance is less than the 5th percentile of pixel-to-centroid distances within that cell.

**Figure 4:**
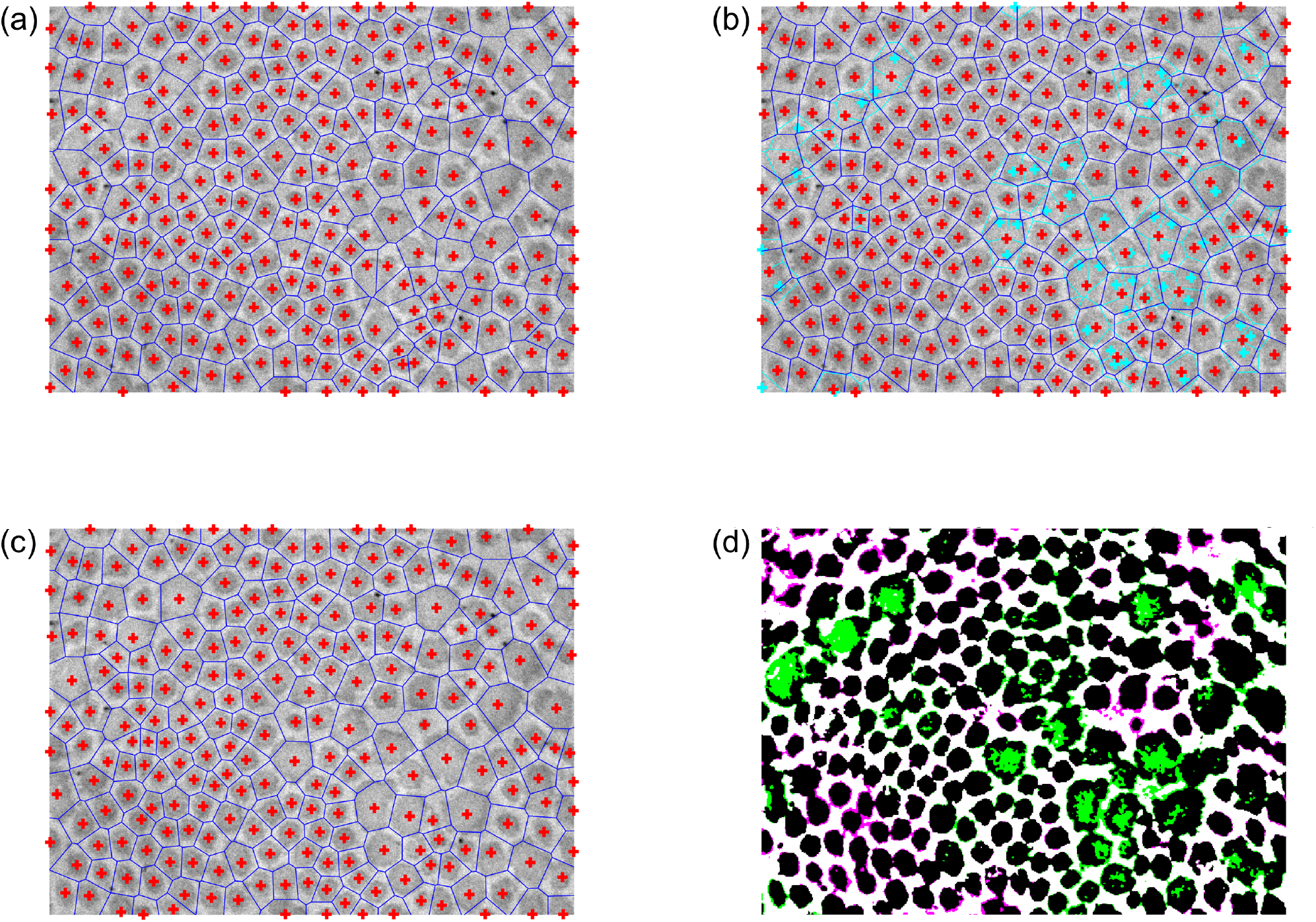
Fibril neighborhoods based on Voronoi tesselation. (a) Initial guess for the fibril neighborhood boundaries (blue) using centroids (red crosses) based on the local peaks in the optimally smoothed *D*_filt_ (*σ*\*). (b) Subsequent interactive correction of the centroid locations. New centroids and boundaries are shown in red and blue, respectively, while the initial guess centroids and boundaries are now shown in cyan for comparison. (c) The final user-approved centroid locations and corresponding Voronoi tesselation. (d) Improved binarization based on the corrected centroid locations and neighborhood boundaries, using a smooth heterogeneous transformation of the intensity field based on characteristic neighborhood intensities that account for intrafibrillar intensity variations. Regions whose categorization changed from non-fibrillar to fibrillar are colored green, while regions whose categorization changed from fibrillar to non-fibrillar are colored magenta. The categorizations of black and white regions are unchanged (fibrillar and non-fibrillar, respectively).

A smooth interpolant *f*_2_ (*u,v*) of the characteristic intensities (defined at the centroids for the purposes of interpolation) is then computed. To ensure *C*^1^ continuity, a natural neighbor area-based local weighting scheme that is determined directly from the above Voronoi tessellation is used to compute the value of the interpolant at each pixel location [26, 27]. The adjusted image is then readjusted using

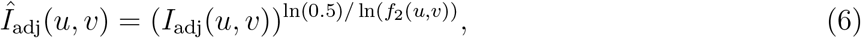

and *Î*_adj_ is binarized (again using Otsu’s method [25]) to yield *Î*_bin_ (Figure 4d). To summarize the overall presence of fibrillar collagen, it is trivial to compute the collagen fibril area fraction as the percentage of *Î*_bin_ consisting of fibrillar pixels.

While the Voronoi tessellation provides a necessary platform from which to refine the image binarization (thereby improving the categorization of fibrillar and non-fibrillar regions), the Voronoi cells do not generally yield a precise delineation of the neighborhood boundaries surrounding each fibril. In particular, when adjacent fibrils differ substantially in size, the shared edge of their Voronoi cells tends to pass through the larger fibril, since the Voronoi tessellation is based only on the fibril centroids (Figure 4c). A more sophisticated approach to determine the true fibril boundaries is thus needed to avoid errors when computing fibril-specific metrics of interest, such as cross-sectional area and aspect ratio. To overcome this limitation of the Voronoi tessellation, the newly segmented fibrillar pixels are therefore clustered by fitting a two-dimensional Gaussian mixture model to the fibrillar pixel coordinates [28]. Note that clustering via the Voronoi tessellation (i.e., *k*-means clustering) is a special case of clustering with a Gaussian mixture model, under the constraint that the component covariance matrices must all be equal and proportional to the identity matrix. Using the Voronoi centroid locations as an initial guess for the *N* cluster means and the Voronoi cell memberships as initial cluster assignments, fitting is performed via an expectation-maximization algorithm [28, 29], wherein computation of the posterior membership probabilities conditional on the current distribution parameters (“E step”) is alternated with an update of the distribution parameters based on the membership probabilities (“M step”). To avoid convergence onto solutions that include rank-deficient component covariance matrices, the component means ***μ**_i_* and proportions *ϕ_i_* are first optimized conditional on the covariance matrices **Σ**_*i*_, then the covariance matrices and proportions are optimized conditional on the means. In practice, an additional benefit of this sequential approach is that it tends to bias the adjusted cluster means to remain close to their initial (i.e., post-Voronoi, user-approved) locations.

After fitting the model, posterior membership probability fields *P_i_*(*u,v*) are computed over the entire image for each cluster (i.e., each fibril) *i*, using

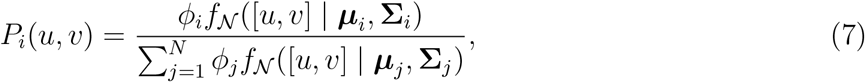

where 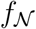 is the normal probability density function. The fibril neighborhood boundaries are then redefined implicitly as *P_i_*(*u,v*) = 0.5 (Figures 5 and 6a). The cross-sectional area of each fibril is computed by summing the areas of all fibrillar pixels within the corresponding neighborhood boundary. At the group level, distributions of fibril area are computed via kernel density estimation of log-transformed area values [30], to enforce a strictly positive support. Under the approximation that fibrillar pixel coordinates belonging to a particular fibril are uniformly distributed within an ellipse, the marginal density of fibrillar pixels along the ellipse’s major axis direction relative to its centroid is

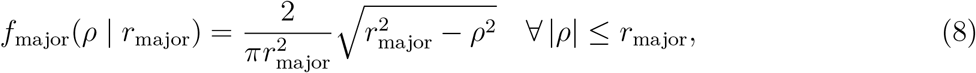

where *r*_major_ is the major radius of the ellipse. Therefore, the variance of fibrillar pixel locations along the major axis is

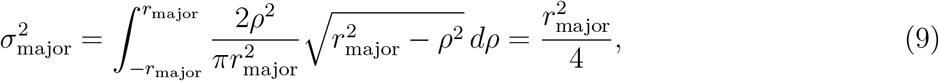

while the variance along the minor axis is analogously 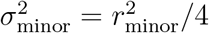. The major and minor radii of the ellipse can thus be computed as *r*_major_ = 2*σ*_major_ and *r*_minor_ = 2*σ*_minor_ respectively [31], where 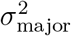 and 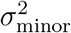 are the maximum and minimum eigenvalues of the covariance matrix for those pixels’ coordinates (Figure 6b). Likewise, the ellipse’s orientation is taken from the eigenvectors of the covariance matrix. As a final correction step, the user may interactively remove any ellipses that fit the actual fibril boundary poorly.

**Figure 5:**
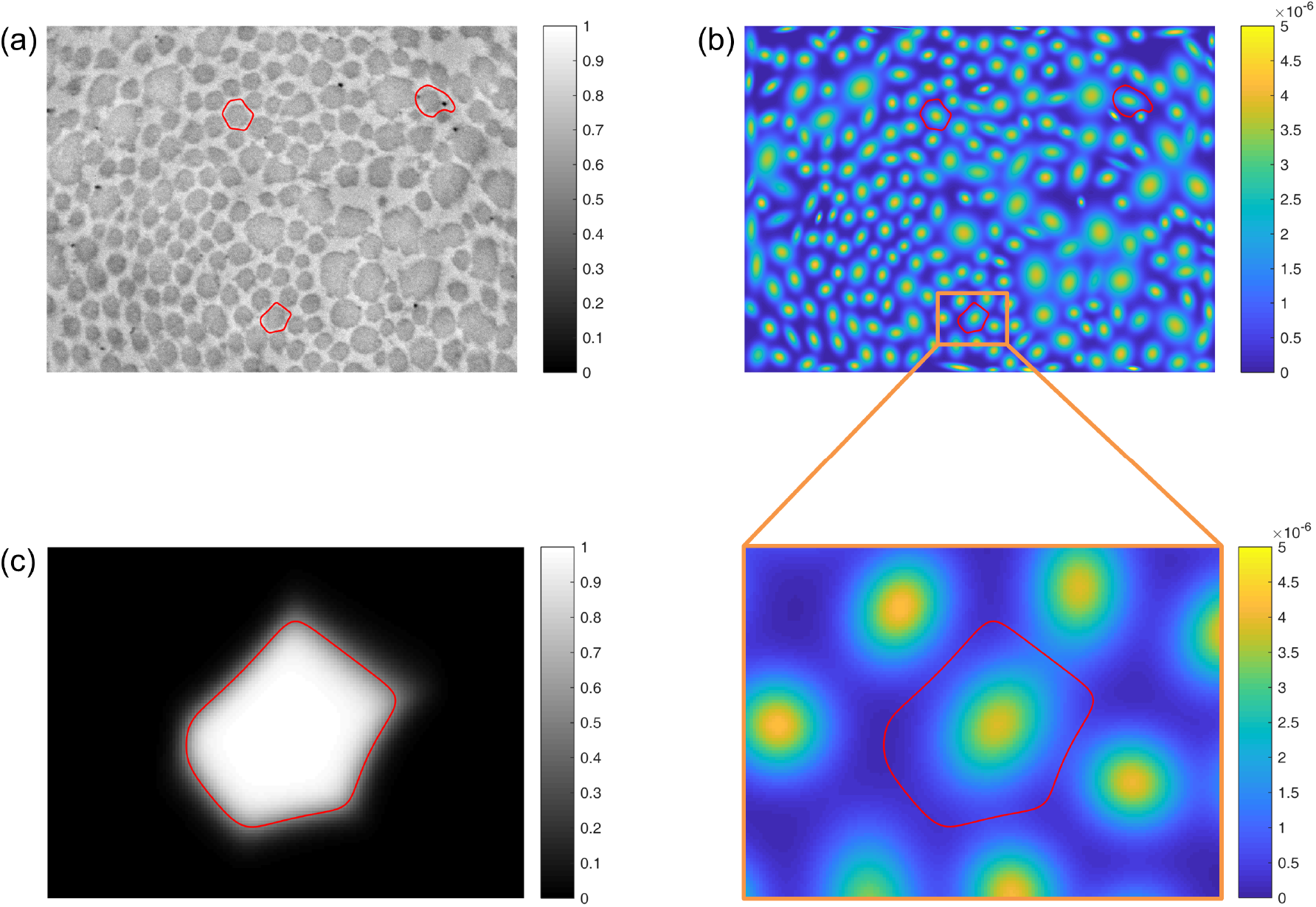
(a) A TEM image, showing improved fibril neighborhood boundaries (red) for three representative fibrils. These boundaries are computed based on fitting an *N*-component Gaussian mixture model to all fibrillar pixel locations. (b) The same three neighborhood boundaries, plotted together with the probability density function of the resultant Gaussian mixture model. A region of interest surrounding one of the representative fibrils is enlarged for clarity. (c) The posterior membership probability field *P_i_*(*u,v*) for the Gaussian component (i.e., fibrillar pixel population) centered in the region of interest. The final neighborhood boundary, defined implicitly as the contour at which *P_i_*(*u,v*) = 0.5, represents that largest *P_i_*-based region that is guaranteed not to overlap with any adjacent fibril neighborhoods, which are defined in the same way.

**Figure 6:**
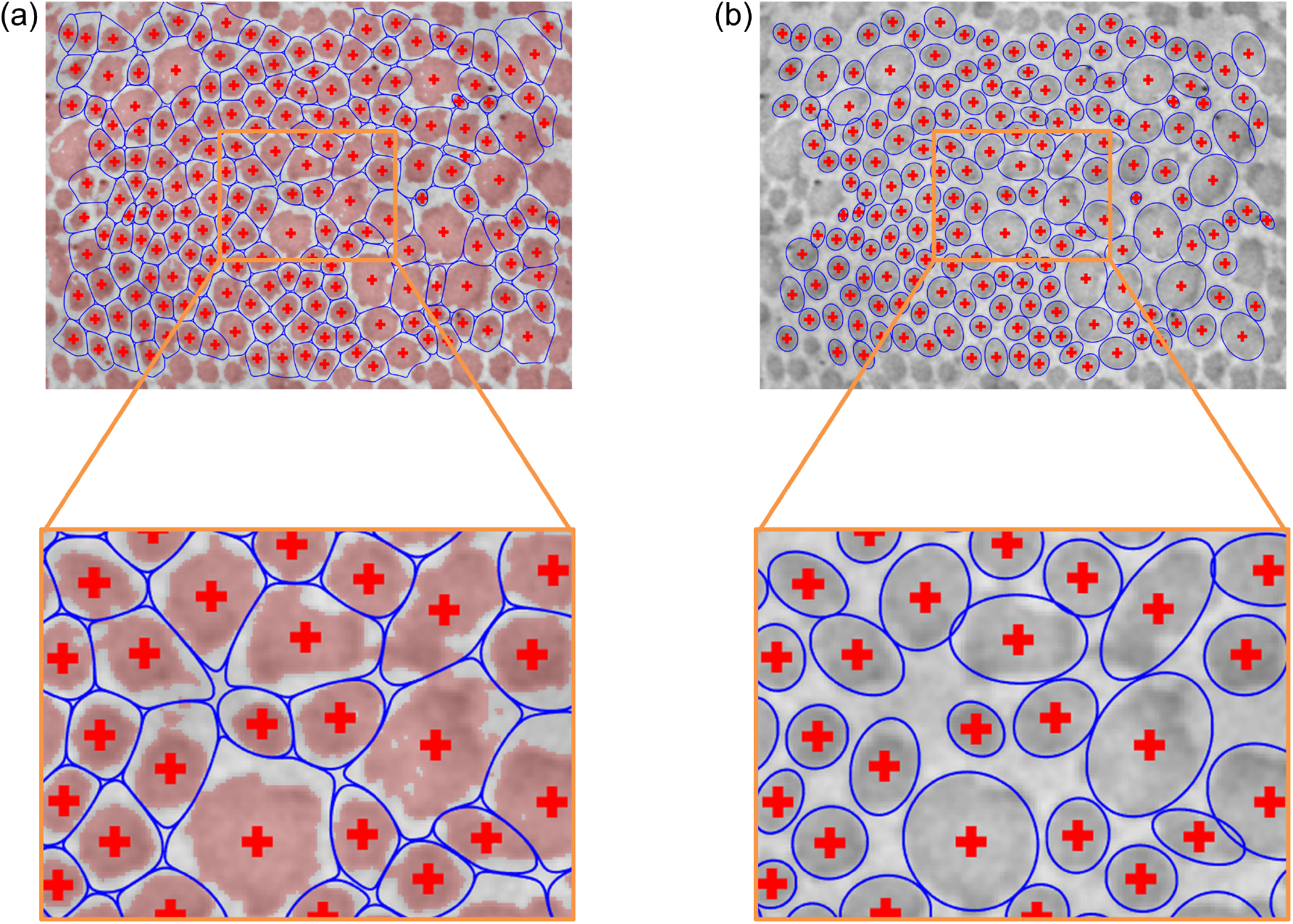
(a) Neighborhood boundaries based on Gaussian mixture modeling for all the fibrils in the image shown. Pixels categorized as fibrillar are overlayed in red. (b) Subsequent approximation of the fibril cross sections, under the simplifying assumption that pixels belonging to each fibril are uniformly distributed within an ellipse.

## 3. ILLUSTRATIVE RESULTS

Our semi-automated TEM image segmentation and analysis pipeline performed well across the entire illustrative data set included in the present study. In particular, we were able to distinguish fibrillar and non-fibrillar regions with minimal user guidance, which enabled efficient extraction of fibril centroid locations and fibril neighborhood boundaries, as well as associated metrics of interest, including fibril cross-sectional area and aspect ratio distributions. Illustrative segmentation and analysis results are shown in Figure 7. Comparing across these images, we found that the different mouse models varied substantially in terms of fibril cross-sectional areas (Figure 8). Specifically, wild-type control fibrils were smaller than both the *Tgfbr1r2* and MFS fibrils, on average. The three area distributions were also substantially different in terms of their variance, with wild-type fibrils being the least variable in size and MFS fibrils being the most variable (Figure 8a). With regard to geometry, the larger fibrils in the *Tgfbr1r2* specimen often had a more irregular shape. Despite this, overall differences in shape as quantified by the fibrils’ aspect ratios were minimal at the population level (Figure 8b). Note that we present these results merely to illustrate our image segmentation and analysis pipeline for a variety of microarchitectures; these findings are not meant to be conclusive or to suggest broader implications.

**Figure 7:**
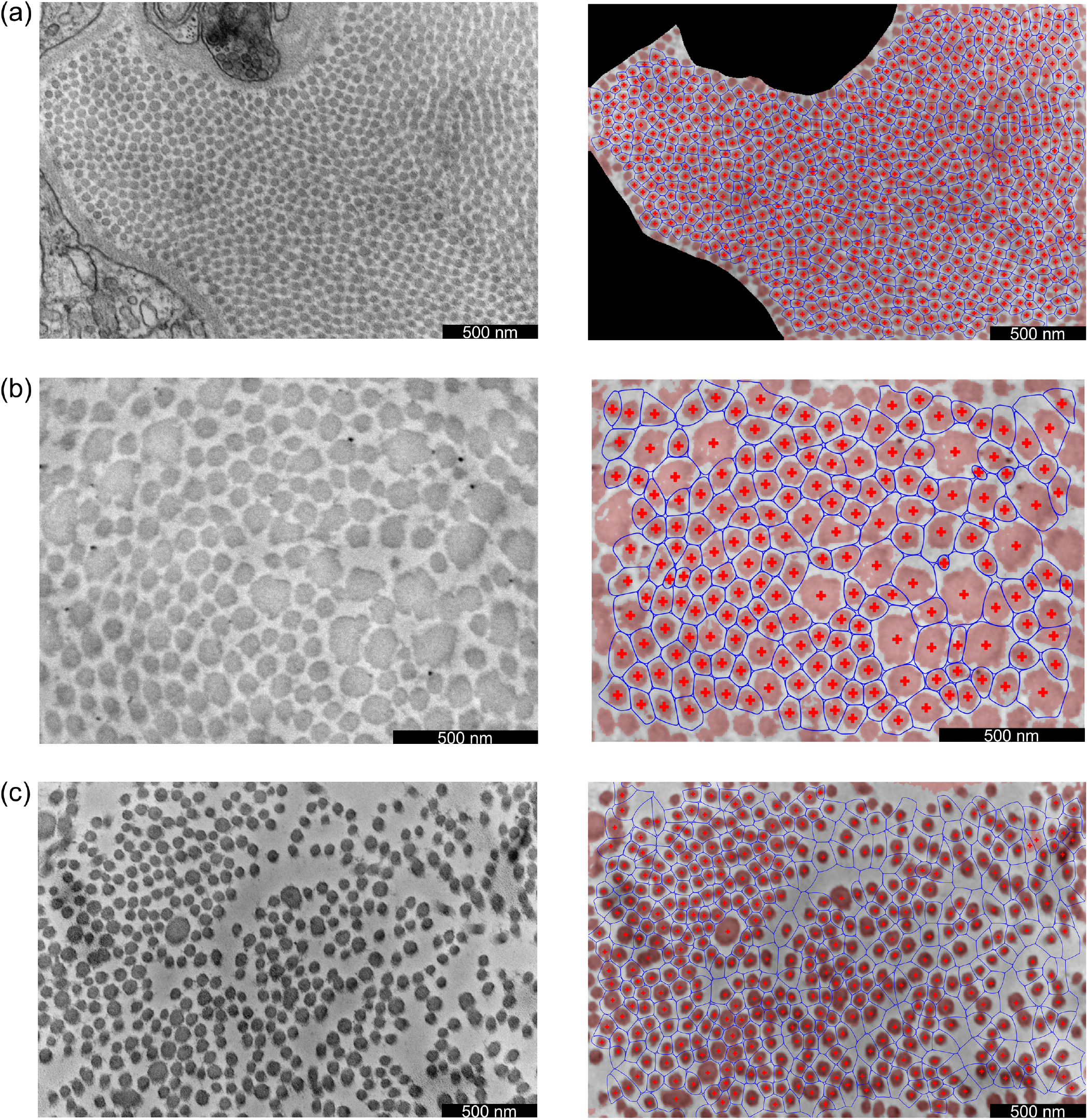
Illustrative results for (a) a wild-type control ATA, (b) a DTA with postnatal disruption of *Tgfbr1r2*, and (c) an ATA representative of MFS (*Fbn1^C1041G/+^*). Acquired TEM images are shown in the left column, while corresponding results are shown in the right column. In the results, pixels categorized as fibrillar have a light red overlay, the fibril centroids are denoted by red crosses, and the fibril neighborhoods are denoted by blue curves.

**Figure 8:**
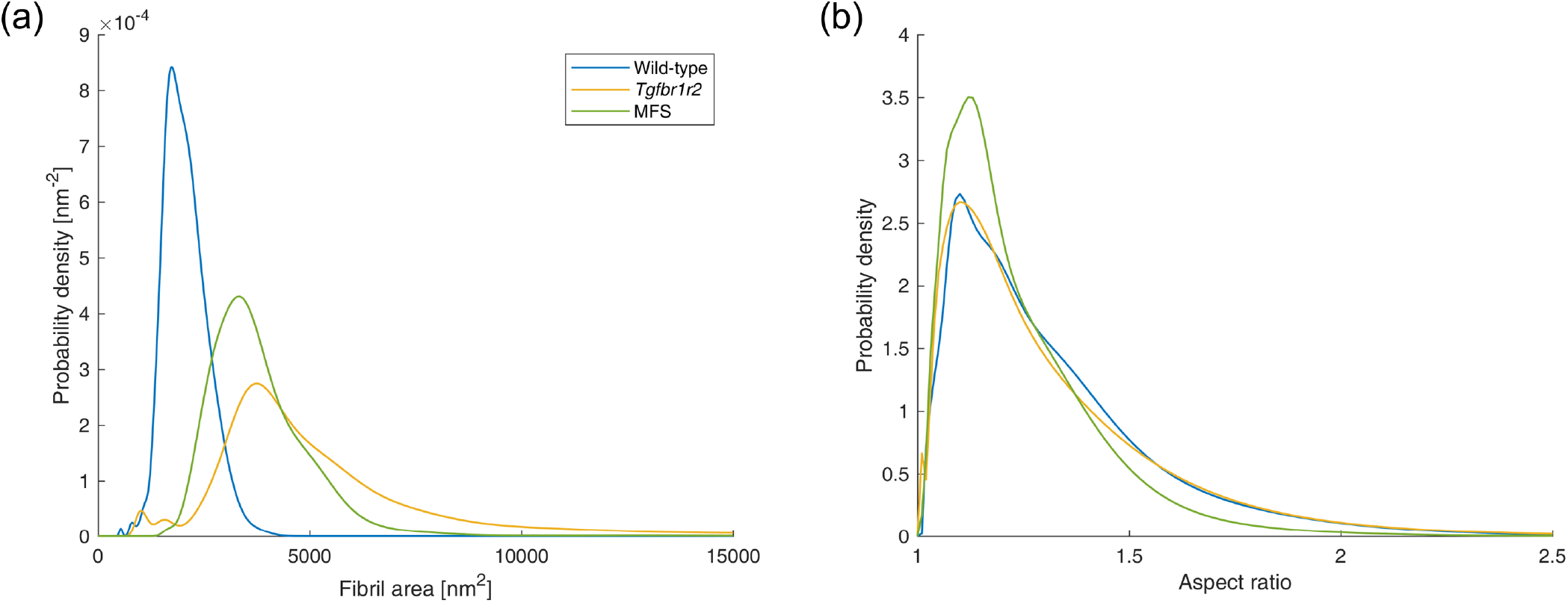
Distributions of (a) fibril cross-sectional area and (b) fibril aspect ratio for the three images shown in Figure 7.

## 4. DISCUSSION

Quantitative analysis of collagen architecture is critical in biomechanical and mechanobiological studies, particularly given the important role of collagen in soft tissue development, homeostasis, adaptation, and disease progression (i.e., via mechanosensing/mechanotransduction and subsequent cellular production and/or removal of tissue constituents). In the present study, we introduce a novel semi-automated pipeline developed to segment individual fibrils from TEM images, which can be used to quickly and robustly quantify geometric characteristics of interest, including fibril crosssectional area and aspect ratio (a measure of circularity). Focusing on vascular applications, we illustrated the pipeline by applying it to images of the thoracic aorta from three different mouse models: wild-type control, *Tgfbr1r2*, and MFS.

Development of this pipeline was motivated by the practical limitations of currently available methods to extract collagen fibril characteristics reliably from TEM images. Since rudimentary thresholding/binarization approaches perform poorly under various circumstances (e.g., Figure 2d), prior studies have relied on interactive tracing of the cross section boundaries for all fibrils in each acquired image [13]. While dependent on tissue characteristics and image magnification, each

acquired TEM image typically contains hundreds of fibrils. Precise interactive tracing therefore requires several hours per image, which has severely limited the sample sizes included in previous multi-group analyses. By contrast, segmentation of fibrils across an entire image can be performed in less than 15 minutes with our novel pipeline, even by a user with no prior training. We found the pipeline to be robust against multiple pathological architectures, namely (1) *Tgfbr1r2*, in which fibril sizes are remarkably variable and larger fibrils exhibit non-uniform image intensity within their cross sections (Figure 7b), and (2) MFS, in which the spatial distribution of fibrils was substantially heterogeneous (Figure 7c). With few exceptions, the segmentation of all individual fibrils was successful, despite several simplifying modeling assumptions utilized in our approach. For example, while the distributions of fibrillar pixels are nearly uniform (rather than Gaussian) within each fibril’s cross section, we found that a Gaussian mixture model performed well for the purposes of defining fibril neighborhood boundaries across the entire image. This high accuracy can be attributed primarily to two factors: (1) although some pairs of fibrils are in direct contact, adjacent fibril boundaries are generally well separated, even in samples where the fibril population density is high (Figure 7a); (2) because the cross sections for the vast majority of fibrils are close to elliptical, their boundaries are well-captured by level sets of their corresponding Gaussian distributions, which are also ellipses.

While the current pipeline has performed well across our validation data set, some limitations remain and should be addressed in future extensions of our approach, particularly in the post-segmentation analysis of the results. Most notably, that aspect ratio distributions were similar across the three specimens (Figure 8b) despite remarkable qualitative differences in fibril crosssectional shape (Figure 7) suggests that there is a need to explore alternative shape comparison metrics, including various measures of deviation between the actual fibril boundary shape and the ellipse approximation [32]. Moreover, while outside the scope of this initial proof-of-concept study, it remains important to apply and assess the pipeline presented herein to substantially larger image sets, which in practice are acquired within application studies focused on specific pathologies [23]. Future evaluation should also be performed with non-vascular samples to examine the broader applicability of this pipeline, and to modify it as needed for different tissues.

From a scientific and clinical perspective, there remains a pressing need to explore how collagen fibril architecture differs across various physiological and pathological conditions, as well as to elucidate the mechanisms through which collagen fibril architecture impacts tissue development, maintenance, and adaptation. While it is well established that fibril architecture contributes significantly to the mechanical behavior of vessel walls [10–13], much remains unknown about the ways in which this architecture influences mechanosensing and mechanotransduction by tissue-resident cells [18, 21]. These processes are the ultimate drivers of growth and remodeling within the extracellular matrix, including the collagen network itself, in response to (patho)physiological mechanical and biological perturbations. The pipeline presented herein, which enables TEM-based quantitative analyses of fibril architecture to be incorporated more ubiquitously in structural investigations of vessel walls, will facilitate continued progress in these longer-term endeavors.

## 5. Conflicts of Interest

The authors declare no conflicts of interest, financial or otherwise.

## 6. Acknowledgments

This material is based upon work supported by the National Institutes of Health (grant no. U01-HL142518 and P01 HL134605).

## Notes

### Competing Interest Statement

The authors have declared no competing interest.

